# A novel locally c-di-GMP-controlled exopolysaccharide synthase required for N4 phage infection of *E. coli*

**DOI:** 10.1101/2021.10.01.462733

**Authors:** Eike H. Junkermeier, Regine Hengge

**Affiliations:** Institut für Biologie / Mikrobiologie, Humboldt-Universität zu Berlin, 10115 Berlin, Germany

**Keywords:** nucleotide second messenger, c-di-GMP, diguanylate cyclase, DgcJ, glycosyltransferase, enterobacterial common antigen, ManNAc, bacteriophage N4, biofilm

## Abstract

A major target of c-di-GMP signaling is the production of biofilm-associated extracellular polymeric substances (EPS), which in *Escherichia coli* K-12 include amyloid curli fibres, phosphoethanolamine-modified (pEtN-)cellulose and poly-N-acetyl-glucosamine (PGA). However, the characterized c-di-GMP-binding effector systems are largely outnumbered by the 12 diguanylate cyclases (DGCs) and 13 phosphodiesterases (PDEs), which synthetize and degrade c-di-GMP, respectively. *E. coli* possesses a single protein with a potentially c-di-GMP-binding MshEN domain, NfrB, which – together with the outer membrane protein NfrA – is known to serve as a receptor system for phage N4. Here, we show that NfrB not only binds c-di-GMP with high affinity, but as a novel c-di-GMP-controlled glycosyltransferase synthesizes a secreted EPS, which can impede motility and is required as an initial receptor for phage N4 infection. In addition, a systematic screening of the 12 DGCs of *E. coli* K-12 revealed that specifically DgcJ is required for the infection with phage N4 and interacts directly with NfrB. This is in line with local signaling models, where specific DGCs and/or PDEs form protein complexes with particular c-di-GMP effector/target systems. Our findings thus provide further evidence that intracellular signaling pathways, which all use the same diffusible second messenger, can act in parallel in a highly specific manner.

**Importance:** Key findings in model organisms led to the concept of ‘local’ signaling, challenging the dogma of a gradually increasing global intracellular c-di-GMP concentration driving the motile-sessile transition in bacteria. In our current model, bacteria dynamically combine global as well as local signaling modes, in which specific DGCs and/or PDEs team up with effector/target systems in multiprotein complexes. Our present study highlights a novel example of how specificity in c-di-GMP signaling can be achieved by showing NfrB as a novel c-di-GMP binding effector in *E. coli*, which is controlled in a local manner specifically by DgcJ. We further show that NfrB (which was initially found as a part of a receptor system for phage N4) is involved in the production of a novel exopolysaccharide. Finally, our data shine new light on host interaction of phage N4, which uses this exopolysaccharide as an initial receptor for adsorption.

## Introduction

Many key cellular functions in bacteria, ranging from adhesion and biofilm formation to development and virulence, are controlled by the second messenger bis-(3′,5′)-cyclic-di-guanosine-monophosphate (c-di-GMP) (12). Remarkably, the genomes of most bacteria encode a multitude of diguanylate cyclases (DGCs) and phosphodiesterases (PDEs) that synthesize and degrade c-di-GMP, respectively (39). In *Escherichia coli* K-12, most of its 12 DGCs and 13 PDEs are not just expressed, but also active at the same time (11, 42, 47). This multiplicity raised the question of how c-di-GMP signaling can be specific, as all of the c-di-GMP-controlled effector/target systems rely on the same diffusible intracellular second messenger (9). Moreover, knockout mutations in particular single DGCs or PDEs were found to result in strong phenotypes without affecting the strikingly low intracellular c-di-GMP level in *E. coli*, which does not exceed 100 nM even in stationary phase cells, i.e. when the characterized effector/target systems are clearly active (42). These seemingly enigmatic observations could be resolved by a model of ‘local signaling’, in which a master PDE (PdeH) maintains a very low global c-di-GMP pool, while specific DGCs, which are directly and locally associated with specific effector/target systems, can act as local and dynamic c-di-GMP sources to trigger specific responses. Similarly, specific PDEs associated with effector/target systems can act as local sinks of c-di-GMP and thus inhibit the regulatory output (recently reviewed in (8)).

Prototypical examples of such locally c-di-GMP-controlled systems have been examined thoroughly in *E. coli*. For example, cellulose synthesis, modification and secretion by the Bcs machinery is not only c-di-GMP-controlled (27, 49), but depends specifically on the diguanylate cyclase DgcC (YaiC) (2). By being directly localized to the core BcsAB complex via protein-protein interactions, DgcC and PdeK serve as a source and sink of c-di-GMP, respectively, for the c-di-GMP-binding PilZ-domain of the cellulose synthase subunit BcsA (36). In this example, the main function of the protein-protein interactions is to co-localize the source and sink of c-di-GMP to its receptor binding site. In addition, protein-protein interactions between specific DGCs/PDEs and their respective effector/target systems can also assume regulatory functions. Thus, the expression of the biofilm regulator CsgD is controlled by the locally acting DgcE-PdeR-DgcM-MlrA signaling module. In this system, the ‘trigger PDE’ PdeR directly binds and thereby inhibits DgcM and the transcription factor MlrA. When PdeR becomes active as a PDE, i.e. degrades c-di-GMP that is provided specifically by DgcE, this direct inhibition is released with two consequences: DgcM can produce c-di-GMP, which results in a local positive feedback, and the transcription of *csgD* is initiated by the DgcM-MlrA complex (10, 22, 44).

Apart from these characterized systems, the considerable number of 12 DGCs and 13 PDEs of *E. coli* K-12 – most of still unknown function – suggests the existence of additional c-di-GMP-controlled systems. In recent years, various approaches have led to the discovery of novel types of c-di-GMP-binding effector components in other bacteria. Among those, the N-terminal domain of MshE-type ATPases (termed MshEN domain), which are involved in Type IV pilus formation of *Vibrio cholerae* as well as bacterial Type II secretion systems of *Pseudomonas aeruginosa* (13, 37), has been identified as a potent c-di-GMP binding receptor. The MshEN domain binds c-di-GMP in a unique fashion, in which a tandem array of two highly conserved 24-residue motifs (Fig. 1A) undergoes extensive hydrophobic interactions with the dinucleotide (50). Bioinformatic studies revealed that the MshEN domain is a ubiquitous regulatory domain. Apart from being present in ATPases of type II and type IV secretion systems, it also seems to be involved in a variety of other bacterial processes, including two-component signaling, protein phosphorylation, polysaccharide secretion and chemotaxis (3, 50). However, the large majoritiy of these proteins have remained uncharacterized.

**FIG 1.**
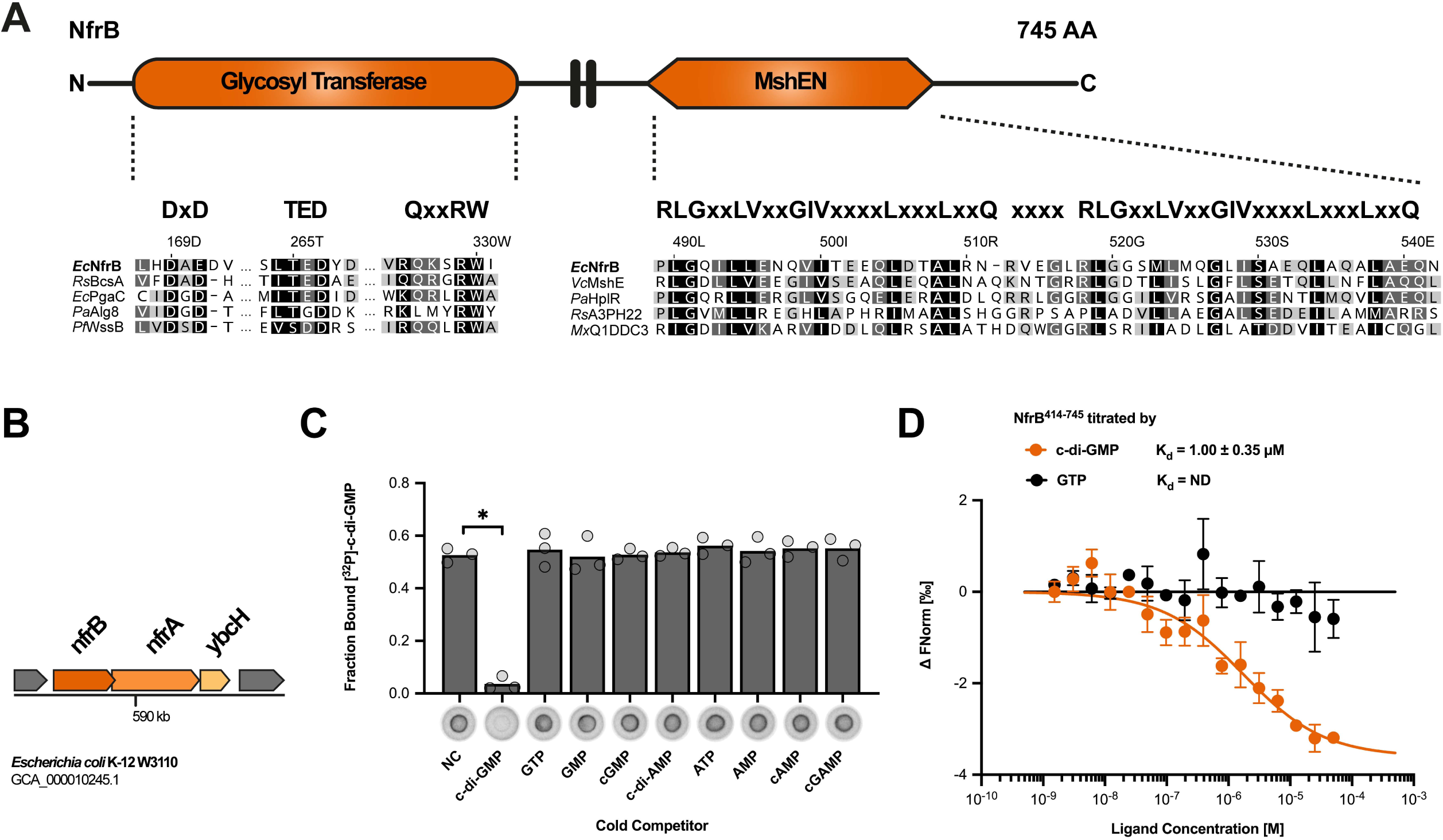
The MshEN domain of NfrB specifically binds c-di-GMP. (A) Domain structure of NfrB (top) and alignment of its glycosyltransferase domain and MshEN domain (bottom). Residues critical for glycosyltransferase activity (20) and binding of c-di-GMP (50) are highlighted above the alignment. (B) Organization of the *nfrBA-ybcH* operon in *E. coli* K-12. The *orf* of *nfrB* is overlapping by 14 nucleotides with the orf of *nfrA*. (C) Differential radial capillary action of ligand assay (DRaCALA) of interactions between purified NfrB^414-745^ (20 μM) incubated with 4 nM [^32^P]-c-di-GMP. Excess (500 μM) of unlabeled mono- and dinucleotides were added to the binding reactions as indicated in competition assays. Protein-ligand mixtures were spotted onto nitrocellulose and allowed to dry before imaging. Individual data points (cycles) and averages (bars) of the calculated fraction bound for three independent experiments are shown. Images of one representative competition assay are shown below the graph. Binding of [^32^P]-c-di-GMP is noticeable by dark spots centered on the nitrocellulose. NC, no competitor. *P* values below 0.001 are marked by a (*) and were determined by a Student *t* test for significant differences compared with the NC control. (D) Interaction of NfrB^414-745^ with c-di-GMP and GTP was measured by microscale thermophoresis (MST). 40 nM of labeled NfrB^414-745^ (RED-NHS dye, NanoTemper Technologies) was incubated with increasing concentrations (0.00153 μM to 50 μM) of c-di-GMP (n=3) or GTP (n=2) and measured by MST using the Monolith NT.115 (NanoTemper Technologies) at 20 % Excitation Power and 40 % MST Power. The change in normalized fluorescence (ΔFnorm in [‰]) is plotted against the concentration of the respective ligand. The dissociation constant (*K*_d_) was quantified by the K_d_ fit of the NanoTemper Analysis software (v2.3).

NfrB is the only protein containing the MshEN domain in *E. coli*. More than 30 years ago, NfrB was found to be part of a receptor system for bacteriophage N4, even though it is located in the inner membrane (16, 17). In addition, phage N4 infection requires the outer membrane protein NfrA, which is directly bound by the phage protein gp65 (16, 17, 23). Moreover, the cytoplasmic protein NfrC (WecB), which also plays a role in the biosynthesis of the enterobacterial common antigen (ECA) has been implicated in phage N4 infection (15, 24).

Here, we show that *E. coli* NfrB is a c-di-GMP binding protein with a novel glycosyltransferase-MshEN domain architecture. We found that its ability to bind c-di-GMP as well as its glycosyltransferase active site are both essential for a successful N4 phage infection. We further provide evidence, that the NfrB-NfrA system produces a novel, yet uncharacterized exopolysaccharide, that not only serves an initial receptor for the phage N4 but that can also impede flagellar activity. Furthermore, successful phage N4 infection is shown to specifically require DgcJ, which directly contacts NfrB by protein-protein interaction, thus establishing a novel locally c-di-GMP-controlled system in *E. coli*. Starting from a very different angle, i.e. phage N4 biology, and using mainly genetics, the accompanying paper came to conclusions that are fully consistent with ours (43).

## Results

### NfrB is a novel c-di-GMP-binding effector protein in E. coli

Since the NfrB protein of *E. coli* K-12 is a larger protein with a fully conserved MshEN domain at its C-terminus (Fig. 1A), we started our study by an analysis of its overall domain structure as well as its genomic context. The N-terminal domain of NfrB shows similarity to family-2 glycosyltransferase (GT) and indeed features the conserved DxD, TED and QxxRW active site signature of processive GTs (20) (Fig. 1A). The two domains of NfrB are linked by two putative transmembrane helices, which most likely anchor NfrB in the inner membrane (Fig. S1A). In *E. coli, nfrB* is encoded in an operon with *nfrA* and *ybcH* (Fig. 1B). A closer inspection of the NfrA protein sequence revealed that NfrA has a classical signal sequence, TPR-rich repeats and a large C-terminal region that most probably forms an outer membrane pore (Fig. S1C). YbcH is a hydrophilic protein with a N-terminal signal sequence-like region that lacks a cleavage site for signal peptidase, i.e. YbcH seems a periplasmic protein, which stays anchored in the inner membrane (Fig. S1F). Taken together, the three proteins NfrB, NfrA and YbcH show the key characteristics of a putative Gram-negative exopolysaccharide synthesis and secretion system, i.e. an inner membrane-located polysaccharide synthase (NfrB), a periplasmatic putative scaffold protein (YbcH) as well as an outer membrane TPR-containing β-barrel protein (NfrA).

Other exopolysaccharide synthesis systems, e.g. cellulose synthase or PGA synthase, are commonly activated by c-di-GMP. NfrB, however, seems the glycosyltransferase, which contains a MshEN domain as a potential c-di-GMP binding domain. Therefore, we purified the soluble MshEN domain of NfrB (NfrB^414-745^) and tested whether it binds c-di-GMP in a radial capillary action of ligand assay (DRaCALA). Specific interaction was indeed observed (Fig. 1C), which was specifically outcompeted by excess unlabeled c-di-GMP, but not by any of the other nucleotides tested (Fig. 1C). To determine the binding affinity, we performed microscale thermophoresis (MST) experiments, which not only confirmed c-di-GMP binding of NfrB^414-745^ but also revealed a K_d_ of 1.0 ± 0.35 μM (Fig. 1D).

### The nfrBA-ybcH operon is post-exponentially expressed, temperature-controlled and co-regulated with flagella

To gain insights into the physiological function of NfrB in *E. coli*, we investigated *nfrB* expression and regulation in *E. coli*. We generated a single copy *nfrB::lacZ* reporter gene fusion, which reflects the promoter activity of *nfrB* (and thus of the entire *nfrBA-ybcH* operon) and monitored its expression during the growth cycle of *E. coli* in liquid LB medium. Overall, *nfrB::lacZ* was expressed at relatively low levels in wild-type cells (Fig. 2A). Expression increased during late exponential phase and again noticeably declined during entry into stationary phase. This pattern was similar, but expression levels were approximately twofold higher at 37°C than at 28°C (Fig. 2A). The decreasing expression of the *nfrB::lacZ* reporter fusion during early stationary phase suggested a negative regulation by the general stress and stationary phase sigma factor RpoS (σ^S^). Indeed, expression of *nfrB::lacZ* remained higher in stationary phase in a *rpoS* mutant background (Fig. 2B).

**FIG 2.**
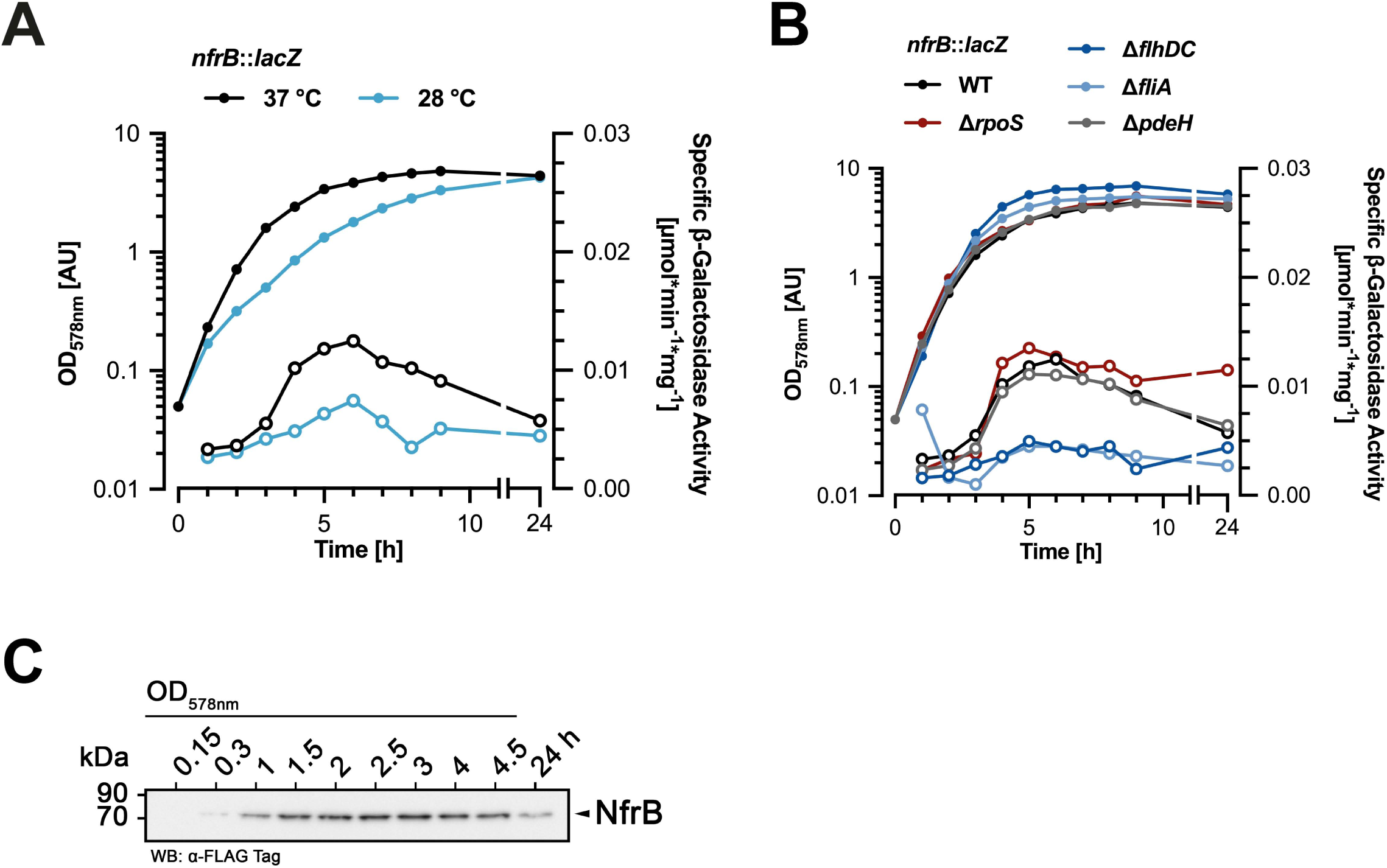
The Nfr system is expressed in post-exponentially growing *E. coli* cells in a temperature-controlled and FliA-activated manner. (A) Expression of the single copy *nfrB*::*lacZ* reporter fusion in the *E. coli* K-12 strain W3110 during the growth in liquid LB medium. OD_578nm_ (closed symbols) and specific β-galactosidase activities (open symbols) were determined during growth at 37°C or 28°C. (B) Expression of the *nfrB*::*lacZ* reporter fusion in W3110 derivatives carrying additional mutations in *rpoS, flhDC, fliA* and *pdeH* was determined as described above for cells growing in liquid LB medium at 37°C. (C) Immunoblot analysis of chromosomally encoded C-terminally 3xFLAG-tagged NfrB in a derivative of strain W3110. Samples were taken at the indicated OD_578_ and after growth for 24 h in liquid LB medium at 37°C.

This transiently increased expression in the late or post-exponential phase is a pattern that is typically found for genes that are regulated by the flagellar control cascade, which involves the flagellar master regulator FlhDC and the flagellar sigma factor FliA (σ^28^). Knocking out either FlhDC or FliA indeed reduced the expression of the *nfrB::lacZ* reporter gene fusion (Fig. 2B). Yet, the absence of FliA also leads to increased intracellular c-di-GMP level (42), since *pdeH*, which encodes the master PDE in *E. coli*, is a FliA-dependent flagellar class 3 gene (6). Therefore, we wanted to rule out the possibility, that c-di-GMP somehow controls the expression of *nfrB*. However, the loss of *pdeH* had no effect on the expression of *nfrB::lacZ* (Fig. 2B). Thus, our data indicate that the *nfrBA-ybcH* operon belongs to the group of flagellar class 3 genes.

In addition, we determined cellular protein levels of NfrB by immunoblot analysis. For this purpose, we inserted a FLAG-tag-encoding sequence close to the 3’ end of *nfrB* in the chromosome. The 3xFLAG-tagged variant of NfrB (NfrB^FLAG^) possesses the tag epitope between A736 and Q737, to avoid any polar effects on the translation initiation of NfrA, as the coding sequences of both genes overlap by 14 basepairs (Fig. 1B). NfrB^FLAG^ showed increasing abundance during postexponential growth of *E. coli*, whereas levels declined again during entry into stationary phase (Fig. 2B). In summary, these results show, that NfrB is predominantly expressed in *E. coli* during post-exponential growth in a FliA-activated manner.

### Disruption of the nfr operon restores the motility defect of a ΔpdeH ΔycgR mutant

The finding that the Nfr system is under control of the flagellar sigma factor FliA – a property that it shares with some other non-flagellar proteins related to c-di-GMP signaling such as PdeH and the PilZ domain protein YcgR – suggested a physiological and possibly regulatory connection between the Nfr system and bacterial motility. Therefore, we tested swimming motility of a Δ*nfrBA-ybcH* mutant strain in semi-solid agar plates but observed no difference compared to the parental strain (Fig. 3A). The loss of the master PDE PdeH renders cells non-motile (6, 30, 40) (Fig. 3A), because the resulting strongly increased intracellular c-di-GMP level (42) activates YcgR, which in its c-di-GMP-bound form functions as a flagellar brake by directly interacting with the flagellar basal body (1, 5). However, knocking out YcgR only partially suppresses this motility defect of a *pdeH* mutant (6), indicating, that an additional factor restrains bacterial motility in the *pdeH ycgR* background. This factor seems to be the Nfr system, since deleting also the *nfrBA-ybcH* operon in addition to *pdeH* and *ycgR* restored wildtype motility (Fig. 3A). Similarly, also single gene disruptions of *nfrB* or *nfrA* could restore full motility of a *pdeH ycgR* mutant (Fig. 3B). Interestingly, however, deleting *ybcH* only partially suppressed the *pdeH ycgR* motility defect (Fig. 3B), suggesting a non-essential role for YbcH in the function of the Nfr system. We conclude that under conditions of increased intracellular c-di-GMP levels (probably required to activate NfrB), the Nfr system can interfere with bacterial motility, most likely by synthesizing and secreting a yet unknown exopolysaccharide.

**FIG 3.**
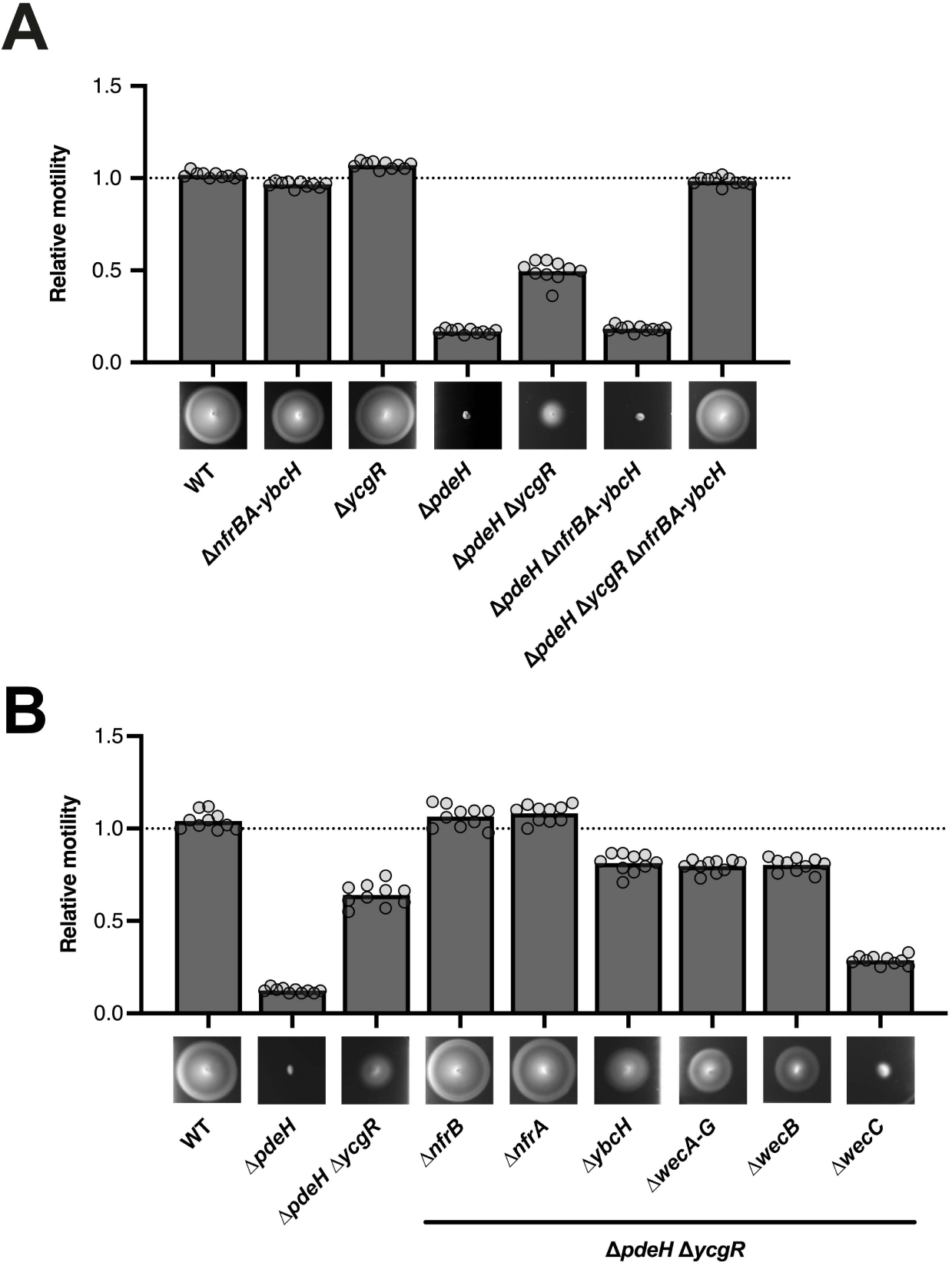
The Nfr system restrains bacterial motility under conditions of elevated intracellular c-di-GMP levels. Motility phenotypes of strain W3110 (WT) and the indicated mutant derivatives were analyzed in TB soft-agar plates (0.3% agar) and quantified after 4.5 h incubation at 37 °C. Representative figures are shown at the bottom of the graphs. The diameters of the motility swarm were measured and normalized to the WT strain. The bar graphs represent the mean of *n* = 10 biologically independent samples. Replicates are shown as individual data points (cycles).

Based on this hypothesis, we reasoned that also a change in substrate availability for the glycosyltransferase activity of NfrB might affect the motility phenotype. For this reason, we further investigated the role of NfrC (WecB), which is also required for phage N4 infection (15) and which was shown to be an epimerase in the production of UDP-N-acetyl-mannosamine (UDP-ManNAc), a precursor for the enterobacterial common antigen (ECA) (24, 34, 41). Indeed, deletion of either the entire *wec* operon (*wecA-G*, encoding all the enzymes required for the biosynthesis of the ECA) or of *nfrC* (*wecB*) alone also showed a moderate suppression of the *pdeH ycgR* phenotype (Fig. 3B). This finding suggests that NfrB uses the product of NfrC (WecB), i.e. UDP-ManNAc, as a substrate for the synthesis of an exopolysaccharide. Interestingly, disrupting *wecC* – the gene next to *wecB* in the *wecA-G* operon – had the opposite effect, i.e. caused an even greater motility defect of the *pdeH ycgR* mutant background (Fig. 3B). Since ECA synthesis is defective in a *wecC* mutant, more UDP-ManNAc produced by the intact NfrC (WecB) in this strain would be available for the synthesis of the exopolysaccharide produced by NfrC.

### Infection of E. coli with bacteriophage N4 requires the activity of the glycosyltransferase domain of NfrB

NfrA, NfrB and NfrC (WecB) have been implicated in the infection of *E. coli* with phage N4 (15, 17). Based on the results described above, we were inspired to gain further insights into the molecular function of the Nfr system by revisiting its role in phage N4 infection. We first analyzed the ability of phage N4 to lyse different derivatives of *E. coli* K-12 W3110 using spot assays. As expected, phage N4 was not able to lyse *E. coli* mutant devoid of NfrB or NfrA, whereas YbcH was dispensable for successful phage infection (Fig. 4A). In addition, a knockout of the biosynthetic pathway of ECA (Δ*wecA-G*) as well as a single gene disruption of *nfrC* (*wecB*) showed a phage N4 plating defect. Remarkably, knocking out WecC, which also results in the absence of ECA, did not change the efficiency of plating of phage N4, indicating that the only contribution of the ECA system to phage N4 infection is the UDP-ManNAc produced by NfrC (WecB). This in turn suggests that the exopolysaccharide produced by NfrB – and not only the NfrA and NfrB proteins *per se* – are involved in phage infection.

**FIG 4.**
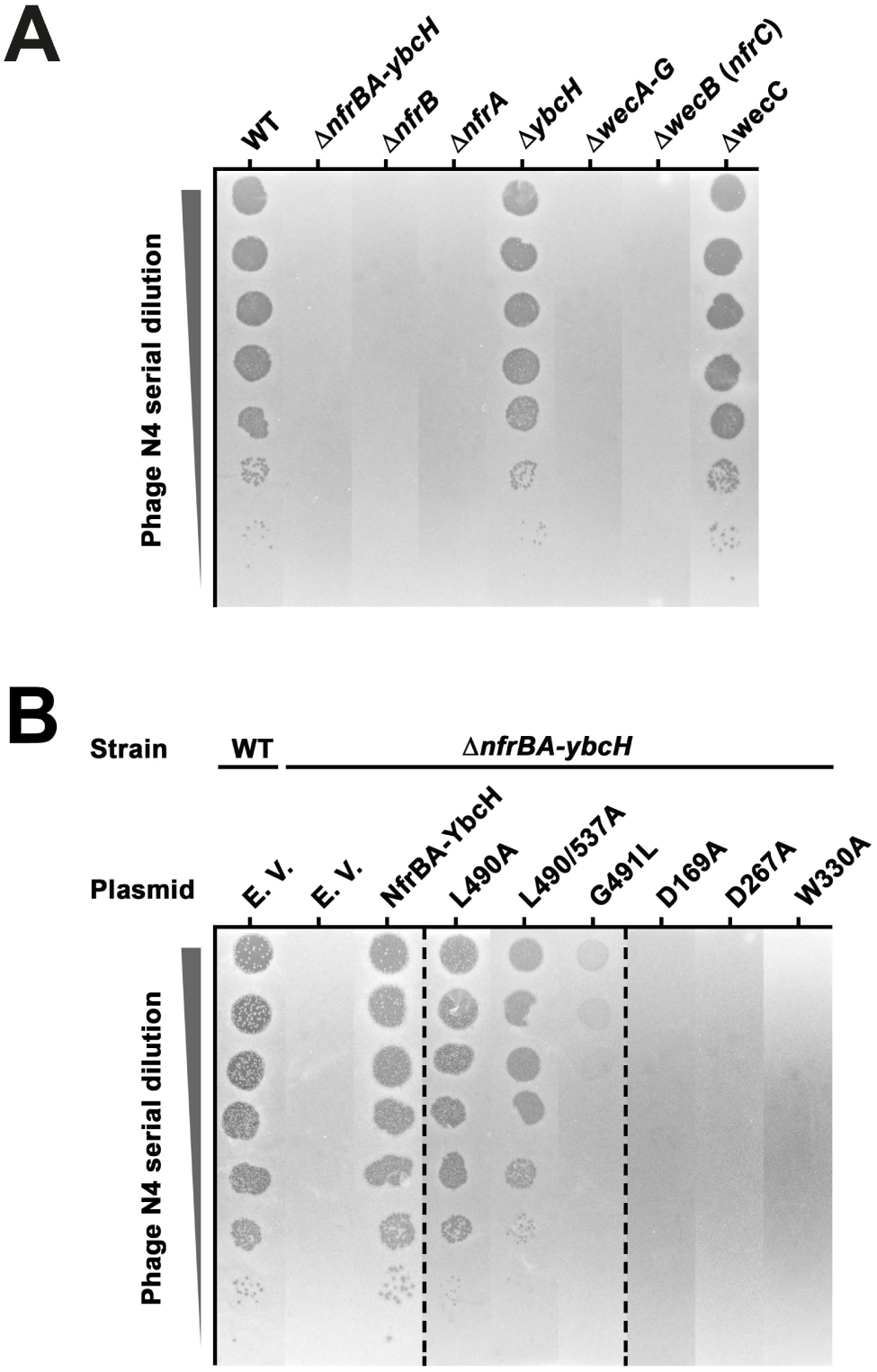
Dissecting the role of the Nfr system in the N4 bacteriophage infection. Plaque formation of phage N4 on *E. coli* K-12 strain W3110 (WT) was tested using serial dilutions (steps of 1:10 dilutions) of a phage N4 lysate spotted (2 μl) on top-agar (LB medium, 1.1 % agar) containing the respective bacterial strains and incubated at 37°C. (A) Plaque formation on strain W3110 and derivatives carrying knockout mutations in the indicated genes. (B) Plaque formation on W3110 (WT) and a derivative strain carrying a deletion removing the entire *nfrBA-ybcH* operon, both transformed with plasmids encoding the wildtype Nfr system (NfrBA-YbcH) or derivatives with the indicated mutations in the MshEN domain (L490A, L490/537A, G491L) or the glycosyltransferase domain (D169A, D267A, W330A) of NfrB. The empty vector (EV) was used as a control.

In order to further dissect the roles of the glycosyltransferase (GT) und MshEN domains of NfrB in phage N4 infection, the *nfrBA-ybcH* operon was cloned onto a low copy number plasmid vector (pAP58), with the *tac* promotor (p_tac_) driving its expression. No inducer (IPTG) was added to the media in order to obtain just moderate (leaky) expression from p_tac_ in our experiments, since the *nfr* operon is expressed at low levels from the chromosome. In addition, we introduced mutations in highly conserved residues of NfrB that are crucial (i) for the glycosyltransferase activity (D169A, D267A and W330A) (20), and (ii) for c-di-GMP binding in the MshEN domain (L490A, L490/537A and G491L) (50). None of these mutations affected the cellular levels of NfrB (Fig. S2). When introduced into a Δ*nfrBA-ybcH* mutant strain, the wild-type construct restored the plating efficiency of phage N4 (Fig. 4B). However, the variants lacking the active sites amino acids of the glycosyltransferase domain of NfrB failed to complement the phage N4 plating defect (Fig 4B). Variants with single (L490A) as well as double (L490/537A) amino acid substitutions in the MshEN domain showed a moderate, but additive reduction in the N4 plaque forming efficiency, whereas the mutation of G491 in the MshEN domain led to the most profound reduction in the N4 plating efficiency (Fig. 4B). Together, these results show, that phage N4 does not only use NfrB and NfrA proteins as the host receptors for infection, but requires both the glycosyltransferase activity of NfrB and its ability to bind c-di-GMP. This indicates that binding of c-di-GMP to the MshEN domain of NfrB allosterically activates its GT domain. This conclusion is in line with the finding reported above, that a mutation specifically in *nfrC* (*wecB*), which eliminates the synthesis of the putative substrate of the GT domain, i.e. UDP-ManNAc, confers resistance to phage N4 infection.

### Specifically DgcJ is required for NfrBA-dependent phage N4 infection and directly interacts with NfrB

One criterium to identify local c-di-GMP signaling is the observation that knocking out distinct DGCs or PDEs leads to highly specific phenotypes (8). Our finding, that the loss of the ability of NfrB to bind of c-di-GMP conferred resistance against the infection with phage N4 (Fig. 4B), raised the question, whether a specific DGC could be required for successful phage infection. Therefore, all the single knockout mutants lacking the 12 active DGCs of *E. coli* K-12 were screened for a phage N4 plating defect (Fig. 5A). In fact, a *dgcJ* deletion showed a severe plating defect, while a *dgcQ* deletion marginally reduced plating efficiency by one order of magnitude. Notably, the *dgcJ dgcQ* double knockout mutant did not show any detectable plaque formation of phage N4 (Fig. 5A). Thus, NfrB seems specifically activated by DgcJ, with DgcQ providing for a minor backup.

**FIG 5.**
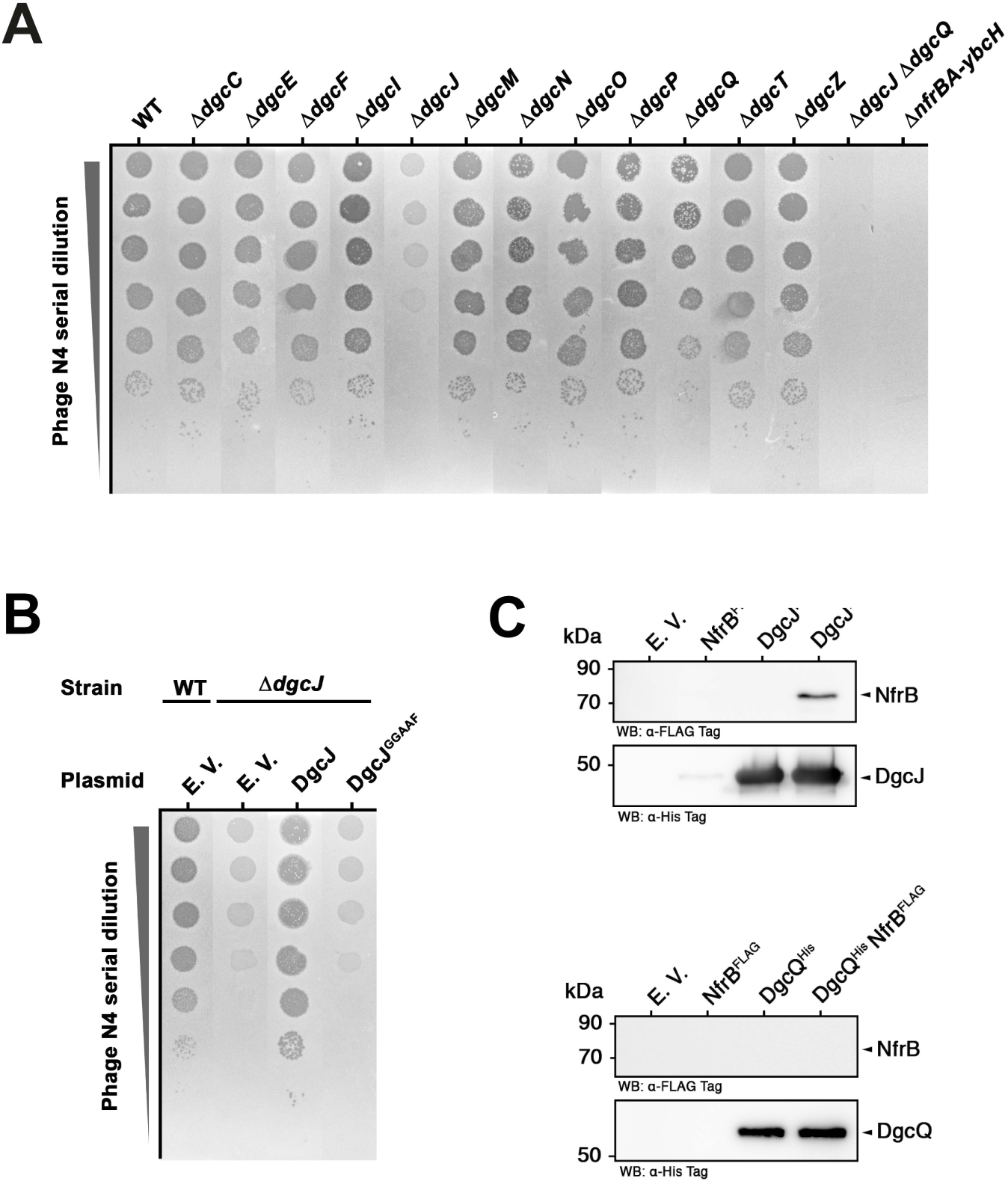
Specifically DgcJ is required for N4 phage infection and directly interacts with NfrB. (A) Plaque formation of phage N4 on strain W3110 (WT) and derivates carrying single deletion mutations in all 12 genes encoding active DGCs, a double deletion of *dgcJ* and *dgcQ* or a deletion of the entire *nfrBA-ybcH* operon was assayed as described in Fig. 4. (B) Plaque formation on a W3110 derivative carrying a deletion in *dgcJ* transformed with a plasmid encoding DgcJ or DgcJ with an active site mutation in its GGDEF domain (DgcJ^GGAAF^). The empty vector (EV) was transformed as a control. (C) NfrB co-purifies with DgcJ. DgcJ^His^ was expressed from the medium copy number vector pRH800 in the presence of NfrB^FLAG^, expressed from the low-copy number plasmid pAP58. Affinity chromatography was performed with the indicated cellular extracts on Ni-NTA resin, which specifically binds the 6xHis epitope of DgcJ^His^. In a similar parallel approach, DgcQ^His^ was purified in the presence of NfrB^FLAG^. Empty vectors were used as controls in combinations as indicated. Eluates were analyzed on SDS polyacrylamide gels, followed by visualization of NfrB (87 kDa), DgcJ (56 kDa) and DgcQ (65 kDa) by immunoblotting using anti-Flag (upper panels) and anti-His6 antibodies (lower panels), respectively.

Next, we focused on the role of DgcJ in phage N4 infection. Therefore, the *dgcJ* gene was cloned on a medium copy number plasmid (pRH800), with p_tac_ driving its expression. No inducer (IPTG) was added to the media in order to not drastically overproduce DgcJ. In addition, we constructed a derivative with active site (A-site) mutations (DgcJ^GGAAF^), to eliminate the DGC activity of the GGDEF domain of DgcJ. When introduced into the *dgcJ* deletion mutant, wild-type DgcJ was able to complement the phage N4 plating defect, whereas DgcJ^GGAAF^ failed to do so (Fig. 5B). These data show, that the N4 infection does not simply require the presence of the inner membrane protein DgcJ, but more specifically the production of c-di-GMP by DgcJ. Therefore, DgcJ is involved in a highly specific c-di-GMP-mediated activation of NfrB.

Specific signaling of a distinct DGC (or PDE) to a particular effector/target system can be expected to occur via a direct protein-protein interaction (8). To test for such interaction, we added a C-terminal 6xHis tag to DgcJ (DgcJ^His^) expressed from pRH800. In parallel, a similar construct on the same vector was obtained with DgcQ (DgcQ^His^), which had shown a minor backup activation of NfrB (Fig. 5A). The coding sequence for NfrB^FLAG^ was cloned together with *nfrA* and *ybcH* on the compatible low-copy number vector pAP58, which allows co-transformation and co-expression of NfrB^FLAG^ and DgcJ^His^ (or DgcQ^His^). All of these proteins were expressed and, when co-expressed, did not affect each others level of expression (Fig. S3). This allowed affinity chromatography or ‘pulldown’ experiments, where DgcJ^His^ or DgcQ^His^ are bound and eluted from a nickel-charged affinity (Ni-NTA) resin (Fig. 3C). When NfrB^FLAG^ was co-expressed with DgcJ^His^, it indeed co-eluted with DgcJ^His^ (Fig 3C). This NfrB^FLAG^/DgcJ^His^ interaction was highly specific, since NfrB^FLAG^ alone was not retained by the Ni-NTA resin and did not co-purify with DgcQ^His^ (Fig. 3C).

In conclusion, among all DGCs of *E. coli*, it is specifically DgcJ that is required for NfrB-dependent infection with phage N4. This role of DgcJ involves its ability to synthesize c-di-GMP. The activation of NfrB by this locally produced c-di-GMP is supported by the direct and specific protein-protein interaction between NfrB and DgcJ.

### NfrB is locally and specifically activated by DgcJ even though DgcQ and DgcE are active in parallel

In principle, our finding that the activation of NfrB depends specifically on the diguanylate cyclase activity of DgcJ would also be compatible with the possibility, that DgcJ may be the only active DGC under the conditions tested (i.e., that the contribution of other DGCs to the global intracellular concentration of c-di-GMP might be negligible). Hence, we addressed the question, whether other DGCs are active during vegetative growth at 37 °C and thus drive up c-di-GMP levels in the *pdeH* mutant, which – via YcgR – interferes with motility.

To test this, we examined whether eliminating other DGCs could suppress the *pdeH* motility defect. When knocked out alone, only the *dgcJ* deletion could relieve the motility defect of the *pdeH* mutant to some extent, whereas deleting either *dgcQ* or *dgcE* had no effect (Fig. 6). However, when, in addition to *dgcJ*, also *dgcQ* or *dgcE* where knocked out, additive effects were observed. Eliminating DgcJ, DgcQ and DgcE all together fully restored the motility of the *pdeH* mutant (Fig. 6). These results indicate, that in vegetatively growing cells at 37 °C, DgcJ, DgcQ and DgcE are all active and contribute to a global pool of c-di-GMP, which in the absence of the master PDE PdeH becomes high enough to inhibit motility via YcgR. However, under conditions where PdeH is present to constantly drain the cellular c-di-GMP pool (42), which allows for motility as YcgR is not activated, it is only DgcJ, which can specifically activate NfrB by direct interaction (Fig. 5C) and thereby allow phage N4 infection.

**FIG 6.**
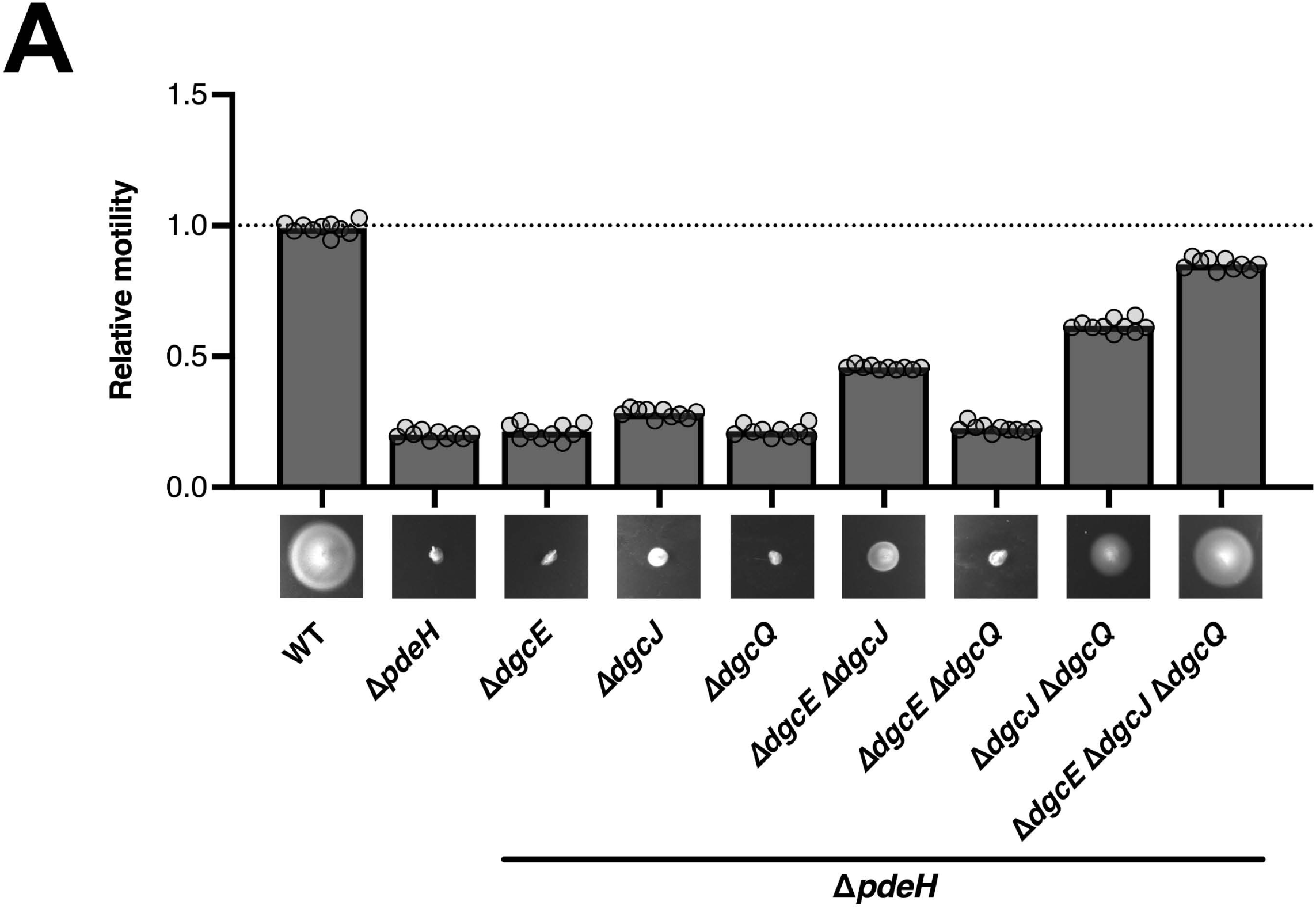
DgcJ, DgcQ and DgcE all contribute to inhibiting motility of a *pdeH* mutant. Motility phenotypes of strain W3110 (WT) and derivatives carrying the indicated mutations in *pdeH, dgcE, dgcJ* and *dgcQ* were analyzed in TB soft-agar plates (0.3% agar) and quantified after 4.5 h incubation at 37 °C. Representative figures are shown at the bottom of the graphs. The diameters of the motility swarms were measured and normalized to the WT strain. The bar graphs represent the mean of *n* = 10 biologically independent samples. Replicates are shown as individual data points (cycles).

### Phylogenetic analysis of the NfrBA-YbcH system shows its frequent genetic linkage to NfrC (WecB)-like enzymes and DGCs

Finally, we analyzed the phylogenetic distribution of NfrB homologs in various bacterial clades with a special focus on the genomic neighbourhoods of the respective genes using TBLASTN searches. 1841 *Ec*NfrB homologs, of which 1101 (60 %) were encoded in different *Escherichia coli* strains, while the remaining ones were present in 406 taxonomically different bacterial species (Fig. 7A). *Ec*NfrB homologs can be found predominantly in γ- and β-proteobacteria, but occasionally also occur in α-proteobacteria and the δ/ε-subdivisions of proteobacteria. All of the identified homologs showed the conserved DxD, TED and QxxRW active site motif in the GT domain (Fig. 7C) and the large majority of 1705 homologs also featured a conserved MshEN domain.

**FIG 7.**
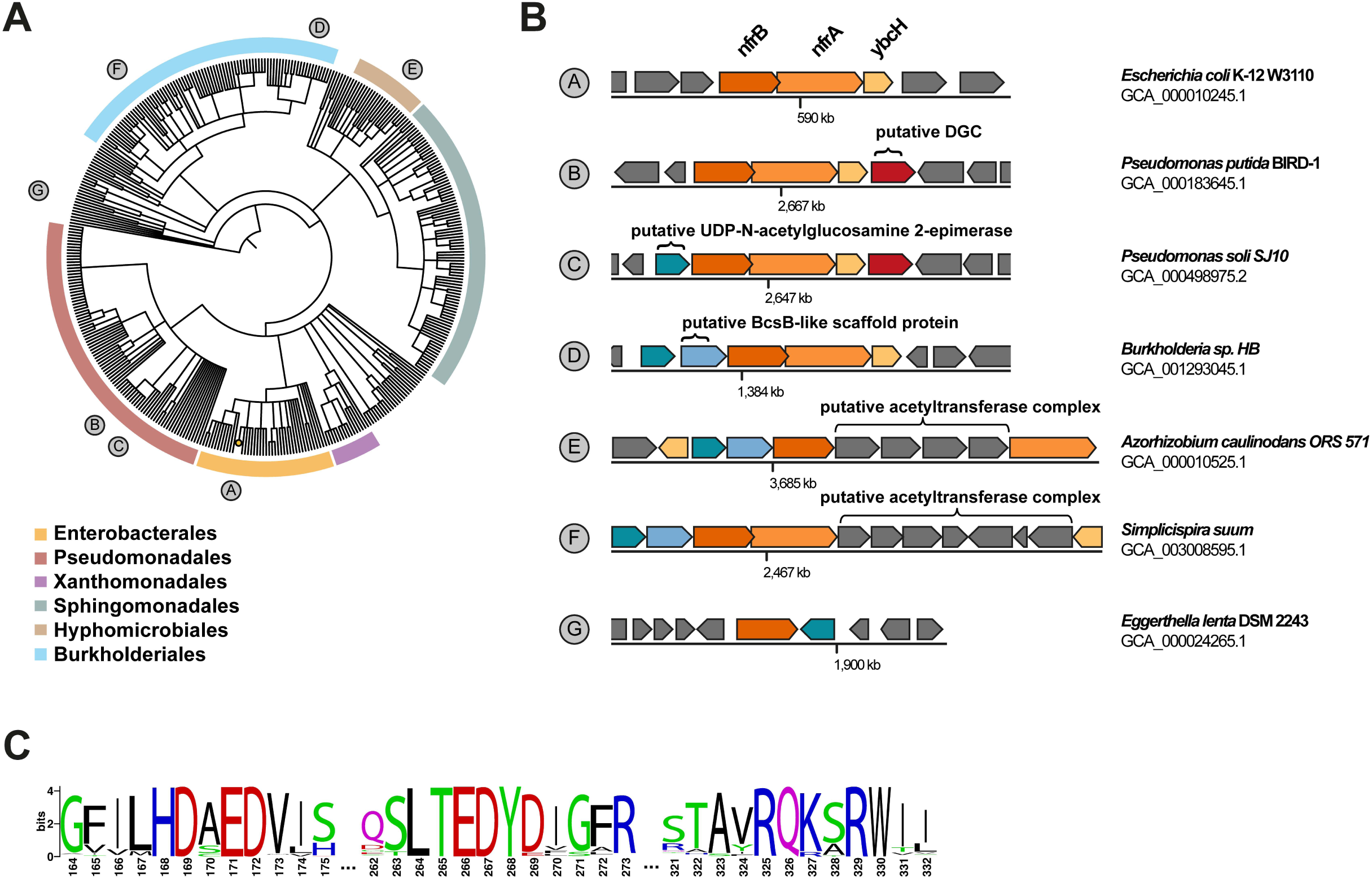
Genomic context and NfrB homologs in various bacterial species. (A) A phylogenetic tree based on the NCBI taxonomy of 406 bacterial species encoding NfrB homologes was generated by phyloT (http://itol.embl.de/) and visualized with iTOL (version 6.3.2.). Bacterial orders are highlighted by the indicated colors. (B) Schematic representations of operons encoding NfrB homologs and their respective genetic background of representative strains, whose position in the phylogenetic tree is shown in (A). (C) Sequence Logo of the 1842 NfrB homologs identified in this study showing the highly conserved active site features of the glycosyltransferase domains. The numbering is according to the corresponding amino acids in NfrB of *E. coli* (see Fig. 1A). The Logo was generated with the alignment in Dataset_S2.

Strikingly, in all organisms identified, which do not encode at least one NfrC (WecB) homolog (i.e. the UDP-N-acetylglucosamine 2-epimerase) in a different biosynthetic pathway (such as the ECA pathway in *E. coli*), a gene encoding a NfrC (WecB) homolog can be found directly associated with the respective NfrB coding sequence (for example in *Pseudomonas putida* BIRD-1 GCA_0000183645.1) (Fig 7B). Moreover, in most pseudomonads, as exemplified by *Pseudomonas soli* SJ10 (GCA_000498975.2), a GGDEF domain protein, i.e. a putative DGC, can be found associated with the *nfr* operon (Fig. 7B). In some cases, e.g. in *Simplicispira suum* (GCA_003008595.1), we identified an additional cluster of genes integrated into the *nfr* gene cluster, which seems related to the acetyltransferase complex that modifies the exopolysaccharide alginate in *Pseudomonas aeruginosa* (26). In rare cases, NfrB homologs can also be found in Gram-positive bacteria like *Eggerthella lenta* DSM 2243 (GCA_000024265.1), which evidently lacks the outer membrane pore (NfrA).

Taken together, our analysis of the local genomic associations of genes encoding NfrB homologs provides further evidence, that the system uses UDP-ManNAc as a substrate to produce an exopolysaccharide. In some cases, this exopolysaccharide may be even modified by an acetyltransferase machinery. The presence of a putative DGC gene immediately downstream, i.e. potentially as a fourth gene in a full *nfrBA-ybcH* operon in some *Pseudomonas* ssp. suggests a specific role of the respective DGC for the Nfr system in these bacteria.

## Discussion

### NfrB is a novel c-di-GMP-binding effector component locally controlled by DgcJ

The highly conserved MshEN domain of NfrB (Fig. 1A) was a strong indication, that NfrB represents a novel c-di-GMP binding effector in *E. coli*. NfrB indeed binds c-GMP specifically (Fig. 1C) with a K_d_ of 1 ± 0.35 μM (Fig. 1D). Thus, NfrB has an affinity in the same range as that of other c-di-GMP effectors in *E. coli*, as exemplified by the K_d_s of 0.84 μM for YcgR (40) or 8.2 μM for BcsA (33). Due to the activity of the strongly expressed ‘master’ PDE PdeH, *E. coli* maintains remarkably low intracellular c-di-GMP levels, ranging from as low as 40-50 nM in vegetatively growing cells (OD_578nm_ of 1) to approximately 80-100 nM during the transition into stationary phase (OD_578nm_ of 3) (42). With a K_d_ that is at least 10-fold higher, NfrB should thus mainly be in the c-di-GMP-free (hence inactive) state under these conditions – if it responds just to the global intracellular c-di-GMP concentration. However, the finding that NfrB is active under these conditions – as demonstrated by the ability of phage N4 to infect *E. coli* in a Nfr-dependent manner (Fig. 4) – suggests local activation by c-di-GMP.

Such an active output of a c-di-GMP-controlled process at global cellular c-di-GMP levels severalfold below the K_d_ of the relevant c-di-GMP-binding effector is one of several criteria that should all be met to unequivocally establish a case of local c-di-GMP signaling (8). Another criterium consist in direct interactions between the specific DGG (and/or PDE) and effector/target component(s) in a signaling protein complex. Such physical vicinity increases the probability of c-di-GMP produced by the specific DGC to either hit the effector binding site or that of a co-localized PDE (8, 36). DgcJ and NfrB were indeed found to directly interact (Fig. 5C). The inability of an A-site point mutation in DgcJ (DgcJ^GGAAF^) to complement the Δ*dgcJ* phenotype (Fig. 5B) indicates that DgcJ functions to provide c-di-GMP locally, so it has a high chance of hitting the MshEN domain of NfrB, resulting in the activation of its GT domain (Fig. 8). In other words, the direct interaction between DgcJ and NfrB plays a scaffolding role similar to DgcC serving BcsA (36), i.e. serves to establish close proximity, rather than also having a direct regulatory impact.

**FIG 8.**
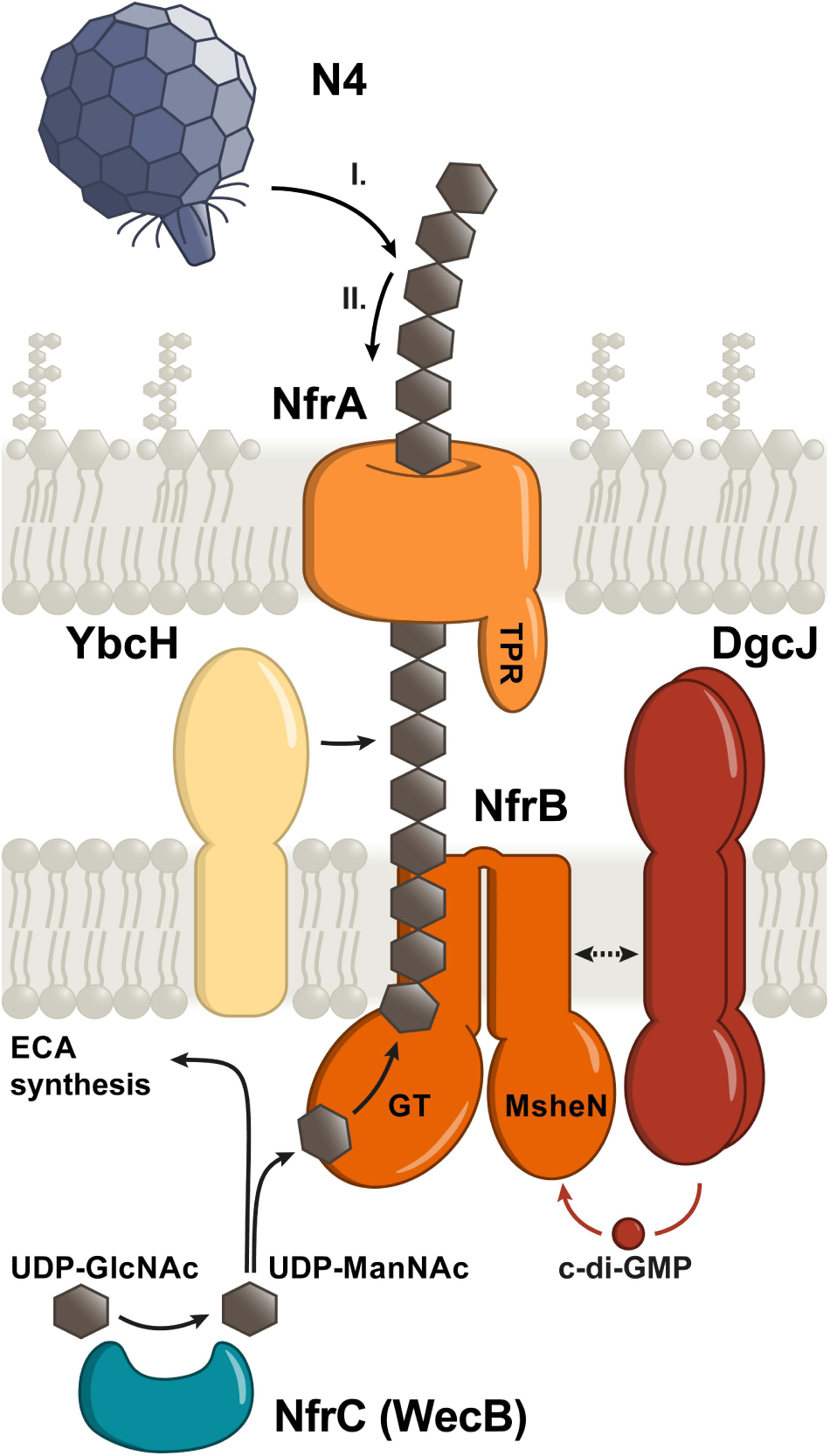
Model of the Nfr/DgcJ system and its role in locally c-di-GMP-activated exopolysaccharide production and bacteriophage N4 adsorption. DgcJ and NfrB co-localize via a direct protein-protein interaction. The C-terminal MshEN domain of NfrB binds c-di-GMP specifically produced by DgcJ, leading to an allosteric activation of the N-terminal glycosyltransferase domain of NfrB. WecB converts UDP-GlcNAc into UDP-ManNAc, which is used for the biosynthesis of the enterobaterial common antigen (ECA). In addition, the glycosyltransferase domain of NfrB uses UDP-ManNAc as a substrate to produce a putative ManNAc-polymer, which is secreted by the outer membrane protein NfrA. YbcH is a periplasmatic protein, which may play an auxiliary role, but is not essential for polysaccharide secretion. Phage N4 binds the exopolysaccharide secreted by the Nfr system as an initial receptor (I.) before interacting with NfrA (II.), which leads to the irriversible adsorption of the phage.

The third criterium for local c-di-GMP signaling are specific phenotypes of mutations that eliminate particular DGCs or PDEs, but not of mutations in other DGCs or PDEs that are concomitantly expressed and active (8). Also this criterium is met by the DgcJ/NfrB system, since the ability of phage N4 to infect *E. coli* in a NfrB-mediated manner strongly and specifically depends on the catalytic activity of DgcJ (Fig. 5A and B), even though DgcQ and DgcE are active under the same conditions (in cells growing at 37°C), as evidenced by the additive effects of all these three DGCs on motility in the absence of the master PDE PdeH (Fig. 6). These observations also confirm the previous insight that for c-di-GMP signaling to act locally, a high active and/or abundant master PDE like PdeH is required to maintain a very low global cellular c-di-GMP pool (42). Notably, the *dgcQ* mutation still had a slight effect on the plating efficiency of phage N4 (Fig. 4A). However, DgcQ did not co-elute with NfrB in our experiments (Fig. 5C), indicating that DgcQ is not as specifically involved in the control of NfrB as DgcJ. Yet, due to the common membrane location of DgcQ and NfrB, DgcQ-produced c-di-GMP may still have some probability to reach NfrB before it is eliminated by PdeH. Based on all these criteria for local c-di-GMP signaling being met here, we therefore propose that DgcJ provides a local source of c-di-GMP right next to NfrB and thereby activates the Nfr system specifically even in the presence of additional active DGCs.

In parallel to our work, another study also reported that c-di-GMP is required for the infection with phage N4 (28). In good agreement with our data, it was found, that a *dgcJ* mutant showed stronger fitness in the presence of phage N4, whereas overexpression of six PDEs (PdeO, PdeR, PdeN, PdeL, PdeB, and PdeI) resulted in a resistance phenotype. That only six of the 13 PDEs of *E. coli* K-12 were able to confer a resistance phenotype, may be due to the remaining PDEs being catalytically inactive under the conditions tested or just not present in the Dub-seq library used (Mutalik et al. 2020). However, the finding that an ectopic expression of various PDEs increases resistance against phage N4 is in line with our proposed local signaling model for NfrB, since NfrB and DgcJ form a specific, yet open signaling module. This allows the DgcJ-produced c-di-GMP to either bind to the MshEN domain of NfrB or diffuse into the cytoplasm, where it gets degraded by PdeH. Expressing additional PDEs from a high copy number plasmid (as in (28)) strengthens the global sink for c-di-GMP in these cells, which will shift the binding equilibrium of NfrB to the c-di-GMP-free and therefore inactive state, which results in the reported phage N4 insensitivity.

Finally, the accompanying publication by Sellner et al. (43) shows that in particular PdeL overexpression confers complete phage N4 resistance. With its LuxR-EAL domain architecture, this ‘trigger PDE’ (10) acts a DNA-binding repressor, which down-regulates its own gene in manner that is allosterically controlled by its c-di-GMP binding and PDE activity (35). Thereby, PdeL does not only reduce the cellular c-di-GMP level in a positive feedback loop, but the new work shows that it also targets additional genes, including the *wec* operon, thereby preventing the expression of NfrC (WecB) (43). Thus, c-di-GMP plays a dual regulatory role in the production of the NfrB-synthesized polysaccharide by controlling *wec* operon transcription and thereby the production of the precursor UDP-ManNAc as well as the glycosyltransferase activity of NfrB.

### The NfrBA system is post-exponentially induced, temperature-controlled, coregulated with flagella and can impede motility

The Nfr system is induced during the post-transcriptional phase of the growth cycle and is transcriptionally regulated by the flagellar control cascade consisting of FlhDC and the sigma factor FliA (Fig. 2B). In addition, it is more strongly expressed at 37°C than at 28°C (Fig. 2A). A similar temperature regulation has also been described for *dgcJ* (previously termed*yeaJ*) (47), which directly stimulates NfrB activity (see above). Overall, this regulatory pattern for the synthesis of the Nfr system and its exopolysaccharide product suggests a physiological function for the NfrB-synthesized exopolysaccharide that may be important within the human host, e.g. in providing protection against recognition by the immune system. Interestingly, lasting upregulation upon upshift to 37°C was recently observed also for other flagellar genes (14), suggesting that temperature input into the Nfr system is connected to its transcriptional regulation by the FlhDC-FliA cascade.

The Nfr systems shares its co-regulation with flagella with several other factors that are involved in c-di-GMP signaling in *E. coli*, i.e. the master PDE PdeH, the c-di-GMP-binding effector YcgR (6) and the GTPase system RdcA/RdcB that eventually – upon a decrease in cellular GTP – directly activates DgcE (31). The coregulation with PdeH is vital for a precise DgcJ-specific control of NfrB activity, since local c-di-GMP signaling depends on a low global c-di-GMP level being maintained by PdeH. On the other hand, YcgR, RdcA/RdcB and the Nfr system all share the ability to tune down motility in a c-di-GMP-controlled manner, albeit through different mechanisms. RdcA/RdcB-activated DgcE provides c-di-GMP that allows YcgR to operate as a c-di-GMP-activated brake, which binds to the flagellar basal body and thereby directly inhibits flagellar rotation from inside the cell (1, 5, 6, 30). By contrast, NfrB is a c-di-GMP-activated glycosyltransferase (Fig. 1) which produces an exopolysaccharide that gets secreted via the outer membrane β-barrel protein NfrA (Fig. 8). Along with YcgR, the Nfr system and therefore most likely its still uncharacterized exopolysaccharide reduce motility in a *pdeH* mutant (Fig. 3). Interestingly, this role of the Nfr system seems analogous to the ability of the exopolysaccharide cellulose to restrain motility of a *pdeH ycgR* mutant of *Salmonella* (52). Our BLAST searches revealed that *Salmonella* does not possess a NfrB homolog and the *E. coli* K-12 W3110 strain used in our motility assays does not produce cellulose (45). Thus, the Nfr-synthesized exopolysaccharide could restrain flagellar rotations by means of steric hindrance in a manner similar to that suggested for cellulose for *Salmonella* (52).

### What kind of polysaccharide does NfrB produce?

The observed effect on motility also correlated with the availability of cellular UDP-ManNAc, since a knockout mutation of *nfrC* (*wecB*) suppressed the motility defect, while it was enhanced by deleting *wecC* (Fig. 3B). Both gene products are enzymes involved in the production of the enterobacterial common antigen (ECA). NfrC (WecB) is a UDP-N-acetyl-glucosamine 2-epimerase responsible for the reversible epimerization between UDP-N-acetyl-glucosamine (UDP-GlcNAc) and UDP-N-acetyl-mannosamine (UDP-ManNAc). WecC catalyzes the following step in the biosynthesis of ECA, in which UDP-N-acetyl-mannosaminuronic acid (UDP-ManNAcUA) is synthesized by a dehydrogenation of UDP-ManNAc. Thus, a *nfrC* (*wecB*) mutant lacks UDP-ManNAc, whereas a *wecC* mutant should have higher levels of UDP-ManNAc, since the next step in ECA synthesis is blocked. Based on the observation that (i) *nfrC* (*wecB*) and *wecC* mutations have opposite effects on the motility phenotype (Fig. 3C) and (ii) NfrC is required for phage N4 infection (Fig. 4A) (15), we propose that the GT domain of NfrB uses UDP-ManNAc as a substrate for the polymerization of a polysaccharide. In wild-type cells, NfrB has to compete with WecC for UDP-ManNAc. In a *wecC* mutant, however, higher UDP-ManNAc levels probably increase the rate of production and secretion of the NfrB-synthesized polysaccharide and thereby steric hindrance of flagellar rotation. That NfrB uses UDP-ManNAc as a substrate is also supported by our finding, that every bacterial species that we found to possess a NfrB homolog, but no NfrC (WecB) homolog associated with a separate biosynthetic pathway (such as the ECA pathway), shows direct genomic association of its respective *nfrB* and *nfrC* (*wecB*) coding sequences (Fig. 7).

### The NfrBA-produced polysaccharide serves as the primary receptor for phage N4

The adsorption of tailed phages to their Gram-negative host surface is a stepwise process. It often includes an initial reversible binding of the phage to cell envelope structures, such as surface-exposed glycans or glycosylated structures. As a secondary step, host receptors in the outer membrane are irreversibly bound, which triggers tail contraction and ejection of the phage DNA into the bacterial cell (reviewed in (29)). Theoretically, phage N4 could bind to the ECA as its initial host receptor. However, in various N4 phage infection studies (16, 28) only mutations in *nfrC* (*wecB*) and none in the other genes of the ECA synthetic gene cluster were found aguing against a role of ECA in phage N4 infection. Our data indicate that the Nfr system most likely uses UDP-ManNAc – the enzymatic product of NfrC (WecB) – as a substrate for the production of an exopolysaccharide. Importantly, mutations in the glycosyltransferase domain of NfrB, as well as its c-di-GMP-binding MshEN domain were also found to generate a phage resistance phenotype (Fig. 4B). We therefore propose that phage N4 uses the NfrB-synthesized exopolysaccharide as its initial receptor (Fig. 8). Based on a similar conclusion, the accompanying study proposes NGR (<u>N</u>4 <u>g</u>lycan <u>r</u>eceptor) as a name for this exopolysaccharide (43). Moreover, quite a low abundance of the Nfr system of three to five copies per cell was reported (16). Consequently, an initial binding to a secreted polysaccharide and subsequent directional movement may be the most efficient way for phage N4 to reach its final host receptor NfrA. Its role as an initial phage receptor also suggests that the exopolysaccharide is not shed from the cells – which could turn it into a dead end trap for the phages – but remains surface-associated.

## Experimental procedures

### Bacterial strains and growth conditions

The strains used in this study are derivatives of *E. coli* K-12 strain W3110 (7). C-terminally 3xFLAG-tagged chromosomally encoded constructs of NfrB (NfrB^FLAG^) were generated by a two-step method similar two the one-step-inactivation protocol (4) as described before (18) using the oligonucleotides listed in Table S1. Knockout mutations in *nfrBA-ybcH, nfrB, nfrA, ybcH, wecA-G, wecB, wecC* are full open reading frame deletions or antibiotic resistance cassette insertions generated by one-step inactivation (4) using the oligonucleotides listed in Table S1. The mutations in all GGDEF/EAL domain-encoding genes as well as in *ycgR, pdeH, flhDC, fliA* and *rpoS* are full *orf* deletion/resistance cassette insertions generated in W3110 and were previously described (30, 42, 47, 51). When required, cassettes were removed as described in (4). P1 transduction (25) was used to transfer the mutations. *E. coli* strain BL21(DE3) (48) was used for protein purification experiments described below. Cells were grown in liquid LB medium under aeration at 28 or 37°C. Antibiotics were added as recommended. Liquid culture growth was followed as optical density at 578 nm (OD 578_nm_).

### Construction of the single copy lacZ reporter fusion

The strain carrying the single copy *nfrB*::*lacZ* reporter fusion also carries a Δ(*lacI-A*)::*scar* deletion as previously described (42, 51). The primers used to construct the fusion are listed in Table S1. PCR fragments were cloned into the *lacZ* fusion vector pJL28, as previously described (51). The fusion was transferred to the att(λ) site of the chromosome via phage λRS45 (46). Single lysogeny was confirmed by PCR (32).

### Bacteriophages and propagation

Phage N4 (GCA_000867865.1) was obtained from the Félix d’Hérelle Reference Center for Bacterial Viruses from the Université Laval (Quebec City, Canada). The phage was propagated on *E. coli* K-12 strains using lysis on plates according to standard protocols (19). Phages were stored and diluted in SM buffer (100 mM NaCl, 8 mM MgSO_4_, 50 mM Tris-Cl) with 0.01 % (w/v) gelatine.

### Protein purification

NfrB^414-745^ was purified as a GST-tagged fusion protein. The coding sequence was cloned on plasmid pGEX-6P-1 (Cytiva 58-9546-48) using the primers listed in Table S1. *E. coli* BL21 Gold strain was transformed with the plasmid and grown to an OD_578nm_ of 0.6 in LB medium at 28 °C, when IPTG (0.1 mM) was added and incubation proceeded for additional 4 h. Cells were harvested and resuspended in lysis buffer (140 mM NaCl, 2.7 mM KCL, 10 mM Na_2_HPO_4_ 1.8 mM KH_2_PO_4_, 5 mM DTT, pH = 7.3) containing protease inhibitor cocktail (complete, EDTA-free; Roche). Cells were disrupted by two passages through a French press. Insoluble material was removed by centrifugation. The supernatant was incubated under gentle shaking overnight with glutathione matrix (Qiagen; 1ml per 1000ml cell culture) at 4°C. The resin was washed with cleavage buffer (50 mM Tris-Cl, 150 mM NaCl, 1 mM EDTA, 1mM DTT, pH = 7.0). PreScission™ protease (Cytiva 27-0843-01; 80 μl protease in 920 μl binding buffer per bed volume) was added and incubated at 4 °C overnight to elute the protein.

Membrane-associated DgcJ^His^ and DgcQ^His^ were purified from IPTG-induced *E. coli* BL21 Gold cells, transformed with pRH800-DgcJ^His^ or pRH800-DgcQ^His^, respectively (pRH800 is a medium copy number p_*tac*_ expression vector). Cells were harvested by centrifugation and resuspended in lysis buffer 50mM Tris (pH8.0), 10 mM MgCl_2_ 300mM NaCl, 1mM EDTA and protease inhibitor cocktail (complete, EDTA-free; Roche). Cells were disrupted by two passages through a French press. Intact cells were removed by centrifugation for 20min at 5000 rpm, total membranes were collected by ultra-centrifugation for 60 min at 36.000 rpm. The membrane pellet was solubilized in 50 mM Tris (pH8.0), 10 mM MgCl_2_ 300 mM NaCl, 5% glycerol, 2% dodecyl-β-d-maltoside (DDM; Roth) for 2h at 4°C. Solubilized and non-solubilized proteins were separated by ultra-centrifugation. The supernatant was incubated with Ni-NTA Agarose (QIAGEN) at 4°C. The resin was washed using solubilization buffer supplemented with 0.05% DDM. Proteins were eluted using solubilization buffer supplemented with 250 mM imidazole.

### Differential Radial Capillary Action of Ligand Assay (DRaCALA)

DRaCALA assayss were performed using 20 μM purified NfrB^414-745^ incubated with 4 nM [^32^P]-c-di-GMP as described (38). Radiolabeled nucleotides were obtained from Hartmann Analytic GmbH. Samples were spotted on nitrocellulose after 10 min incubation at room temperature.

### Microscale Thermophoresis

NfrB^414-745^ was labeled using the Protein Labeling Kit RED-NHS (NanoTemper Technologies). The labeling reaction was performed according to the manufacturer’s instructions in the supplied labeling buffer applying a concentration of 20 μM protein at room temperature for 30 min in the dark. Unreacted dye was removed with the supplied dye removal column equilibrated with MST buffer (137 mM NaCl, 2.7 mM KCl, 10 mM Na_2_HPO_4_, 1.8 mM KH_2_PO_4_, 0.05% Tween). The degree of labeling was determined using UV/VIS spectrophotometry at 650 and 280 nm. The labeled protein was adjusted to 80 nM with MST buffer. c-di-GMP and GTP was dissolved in MST buffer and a series of 16 1:2 dilutions was prepared using the same buffer. For the measurement, each ligand dilution was mixed with one volume of labeled protein, which led to a final concentration of 40 nM and final ligand concentrations ranging from 0.00153 μM to 50 μM. After 10 min incubation, the samples were loaded into Monolith NT.115 Premium Capillaries (NanoTemper Technologies). MST was measured using a Monolith NT.115 instrument (NanoTemper Technologies) at an ambient temperature of 25°C. Instrument parameters were adjusted to 20 % LED power and 40 % MST power. Data of three independently pipetted measurements were analyzed (MO.Affinity Analysis software version 2.3, NanoTemper Technologies).

### Determination of β-galactosidase activity

β-Galactosidase activity was assayed by use of o-nitrophenol galactoside (ONPG) as a substrate and is reported as μmol o-nitrophenol min^−1^ (mg cellular protein)^−1^ (25). Experiments were done at least twice, and a representative experiment is shown. OD_578nm_ was determined and measurements were performed as with cells grown in liquid LB medium.

### Motility assay

Bacterial motility was tested on swim plates containing 0.5 % bacto-tryptone, 0.5 % NaCl and 0.3 % agar. A 3 μl volume of overnight culture (OD adjusted) was inoculated into the swim plates and cells were allowed to grow and swim for 4.5 h at 37°C.

### SDS polyacrylamide gel electrophoresis and immunoblot detection

Proteins were detected by SDS polyacrylamide gel electrophoresis (SDS-PAGE) and immunoblotting as previously described (21) using antibodies against the Flag epitope (Sigma) or the 6xHis-tag (Bethyl Laboratories, Inc.) at 1:10.000 dilution. Anti-rabbit or anti-mouse IgG horseradish-peroxidase conjugate from donkey (GE Healthcare) was used (at 1:20.000 dilution) for protein visualization in the presence of Western Lightning Plus-ECL enhanced chemiluminescence substrate (PerkinElmer). The WesternSure® Pre-stained Chemiluminescent Protein Ladder (Li-cor) was used as a molecular mass standard.

### Identification of NfrB holomogs

NfrB homologs were identified by using the *E. coli* NfrB protein as a querie to perform TBLASTN searches of the NCBI nonredundant protein database. The dataset (dataset_S1) was manually curated (Geneious software, Geneious Prime® 2021.2.2) by removing false-positive hits, as well as disrupted operons (e.g. by mobile genetic elements or mutations in coding sequences). The tree was generated with phyloT (phyloT.biobyte.de) by the NCBI taxonomy of the identified species, in which NfrB homologs were found. iTOL (itol.embl.de) was used to generate the tree. The protein alignment of the 1842 identified homologs was generated with a Genious Alignment (Blosom62 cost matrix, gap open penalty of 12, gap extension penalty of 3 and 2 refinement iterations).

### Additional software tools

Image J (Schneider 2012) was used to calculate the swim diameters of motility plates and intensities of images of phosphorimager films of DRaCALA assays. Sequence logos were generated with the WebLogo service (https://weblogo.berkeley.edu/). Prism 9.2.0 (283) (GraphPad Software, San Diego, California USA) was used for generating graphs and statistical analysis.

## Acknowledgments

We thank Urs Jenal for sharing data prior to publication.

This work was supported by the Deutsche Forschungsgemeinschaft (DFG grants He1556/21-1 and He1556/21-2, as part of DFG Priority Programme 1879 “Nucleotide Second Messenger Signaling in Bacteria”, awarded to RH).

## Author contributions

Concept of the study and design of experiments: EJ, RH; experiments and bioinformatic analyses: EJ; interpretation of experimental data: EJ, RH; writing of the paper: EJ, RH.

