## Supplemental Figures 1-3 and Table 1 for "A novel locally c-di-GMP-controlled exopolysaccharide synthase required for N4 phage infection of *E. coli*"

### Supplementary Figures

**A**

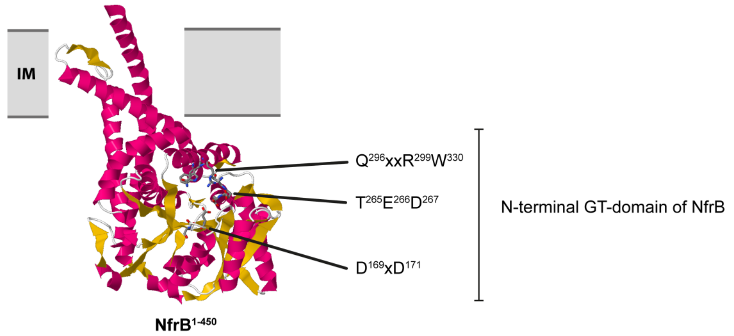


**B**

```
>EcNfrA
MKENNLNRVI GWSGLLLTSL LSTSALADNI GTSAEELGLS DYRHFVIYPR LDKALKAQKN
NDEATAIREF EYIHQQVPDN IPLTLYLAEA YRHFGHDDRA RLLLEDQLKR HPGDARLERS
LAAIPVEVKS VTTVEELLAQ QKACDAAPTL RCRSEVGQNA LRLAQLPVAR AQLNDATFAA
SPEGKTLRTD LLQRAIYLKQ WSQADTLYNE ARQQNTLSAA ERRQWFDVLL AGQLDDRILA
LQSQGIFTDP QSYITYATAL AYRGEKARLQ HYLIENKPLF TTDAQEKSWL YLLSKYSANP
VQALANYTVQ FADNRQYVVG ATLPVLLKEG QYDAAQKLLA TLPANEMLEE RYAVSVATRN
KAEALRLARL LYQQEPANLT RLDQLTWQLM QNEQSREAAD LLLQRYPFQG DARVSQTLMA
RLASLLESHP YLATPAKVAI LSKPLPLAEQ RQWQSQLPGI ADNCPAIVRL LGDMSPSYDA
AAWNRLAKCY RDTLPGVALY AWLQAEQRQP SAWQHRAVAY QAYQVEDYAT ALAAWQKISL
HDMSNEDLLA AANTAQAAGN GAARDRWLQQ AEKRGLGSNA LYWWLHAQRY IPGQPELALN
DLTRSINIAP SANAYVARAT IYRQRHNVPA AVSDLRAALE LEPNNSNTQA ALGYALWDSG
DIAQSREMLE PAHKGLPDDP ALIRQLAYVN QRLDDMPATQ HYARLVIDDI DNQALITPLT
PEQNQQRFNF RRLHEEVGRR WTFSFDSSIG LRSGAMSTAN NNVGGAAPGK SYRSYGQLEA
EYRIGRNMLL EGDLLSVYSR VFADTGENGV MMPVKNPMSG TGLRWKPLRD QIFFIAVEQQ
LPLNGQNGAS DTMLRASASF FNGGKYSDEW HPNGSGWFAQ NLYLDAAQYI RQDIQAWTAD
YRVSWHQKVA NGQTIEPYAH VQDNGYRDKG TQGAQLGGVG VRWNIWTGET HYDAWPHKVS
LGVEYQHTFK AINQRNGERN NAFLTIGVHW
```

**C**

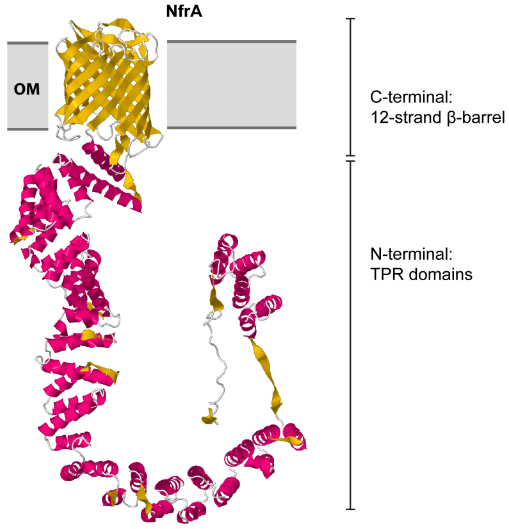


**D**

The sequence name: EcNfrA
Per-protein P-value for being TPR: 1.1E-16
Probability for being TPR: 100.00%

| Repeat | Begin | Alignment | End | P-value |
|---|---|---|---|---|
| TPR | 47 | -- IYPRLDKALKAQKNNDEATAIREFEYIHQCVPDN-- | 80 | 7.9e-04 |
| TPR | 81 | -- IPLTLYLAEAYRHFGHDDRARLLLEDQLKRHPGD-- | 114 | 3.2e-05 |
| TPR | 115 | -- ARLERSLAAIPVEVKSVTTVEELLAQCKACDAAP-- | 148 | 2.8e-01 |
| TPR | 150 | -- LRCRSEVGQNALRLAQLPVARAQLNDATFAASPE-- | 183 | 4.9e-02 |
| TPR | 185 | -- KTLRTDLLQRAIYLKQWSQADTLYNEARQQNTLS-- | 218 | 7.2e-03 |
| TPR | 219 | -- AAERRQWFDVLLAGQLDDRILALCSCGIFTDPQS-- | 252 | 6.4e-02 |
| TPR | 612 | -- ANAYVARATIYRQRHNVPAAVSDLRAALELEPNN-- | 645 | 8.9e-11 |
| TPR | 646 | -- SNTQAALGYALWDSGDIAQSREMLEPAHKGLPDD-- | 679 | 5.2e-06 |
| TPR | 680 | -- PALIRQLAYVNQRLDDMPATQHYARLVIDDIDNQ-- | 713 | 9.5e-03 |

**E**

```
>EcYbcH
MRKFIFVLLT LLLVSPFSFA MKGIIWQPQN RDSQVTDTQW QGLMSQLRLQ GFDTLVLQWT
RYGDAFTQPE QRTLLFKRAA AAQQAGLKLI VGLNADPEFF MHQKQSSAAL ESYLNRLLAA
DLQQARLWSA APGITPDGWY ISAEIDDLNW RSEAARQPLL TWLNNAQRLI SDVSAKPVYI
SSFFAGNMSP DGYRQLLEHV KATGVNVWVQ DGSGVDKLTA EQRERYLQAS ADCQSPAPAS
GVVYELFVAG KGKTFTAKPK PDAEIASLLA KRSSCGKDTL YFSLRYLPVA HGILEY
```

**F**

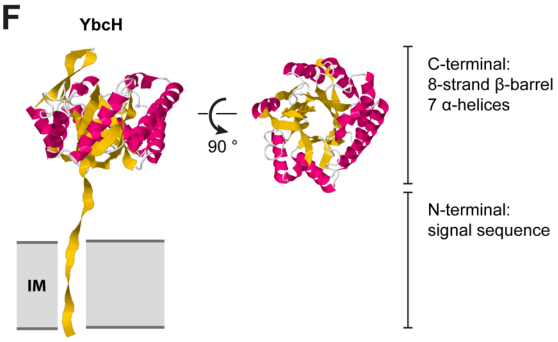

**Figure S1. The Nfr system shows the typical arrangement of a polysaccharide secretion system.**

**A:** Structure prediction of NfrB (identifier AF-P0AFA5-F1) by AlphaFold v2.0 (Jumper , J., R. Evans, A. Pritzel, T. Green, M. Figurnov, O. Ronneberger, et al. 2021. Highly accurate protein structure prediction with AlphaFold. Nature 596:583-589). The N-terminal domain ($NfrB^{1-450}$) is displayed as a cartoon backbone representation. Helical parts are colored magenta, beta sheet structures are colored yellow. Three predicted alpha helices could serve as membrane anchors in the inner membrane. In this configuration, the active site signature (DxD, TED and QxxRW) forms a putative open cavity for substrate binding.

**B:** Amino acid sequence of NfrA (Uniprot identifier P31600). The signal sequence is highlighted in gray and a putative cleavage site for the signal peptidase marked by an arrow head. Predicted TPR-repeats (see D) are shown in green and the predicted C-terminal beta barrel is marked in yellow (see C).

**C:** Structure prediction of NfrA (identifier AF-P31600-F1) by AlphaFold 2.0. A 12-stranded β-Barrel (yellow) could serve as a pore for the secretion of an exopolysaccharide synthetized by NfrB.

**D:** Tetratricopeptide repeat motifs of NfrA as predicted by TPRpred (Gabler, F., S. Z. Nam, S. Till, M. Mirdita, M. Steinegger, J. Söding, A. N. Lupas, and V. Alva. 2020. Protein sequence analysis using the MPI bioinformatics toookit. Curr. Protoc. Bioinformatics **72:**e108; https://toolkit.tuebingen.mpg.de/tools/tprpred).

**E:** YbcH (Uniprot identifier P37325) lacks a cleavage site for the signal peptidase in the signal sequence (gray).

**F:** AlphaFold v2.0. predicts a TIM barrel-like structure (8 strand beta barrel surrounded by 7 α-helixes) for YbcH (AF-P37325-F1). YbcH most likely remains anchored in the inner membrane with its N-terminus.

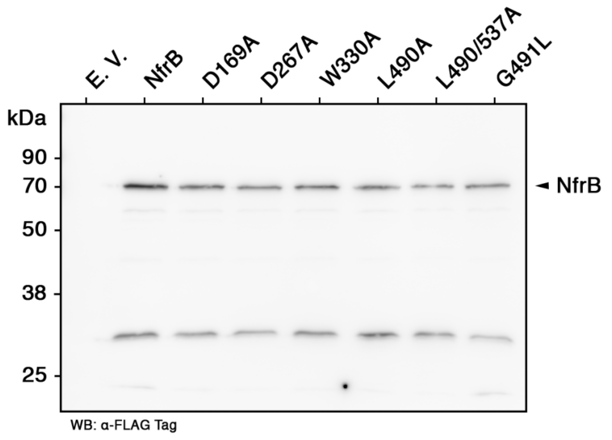


**Figure S2. Single amino acid substitution at relevant positions do not affect cellular levels of NfrB.**

$NfrB^{FLAG}$A-YbcH (with the wildtype sequence as well as with the indicated amino acid substitutions) was expressed from plasmid pAP58 in an *E. coli* K-12 strain W3110 derivative strain carrying a deletion of the *nfrBA-ybcH* operon. Samples were taken from cells grown in liquid LB Medium at 37 °C for 6 hours. The cellular levels of $NfrB^{FLAG}$ were analysed by immunoblot analysis.

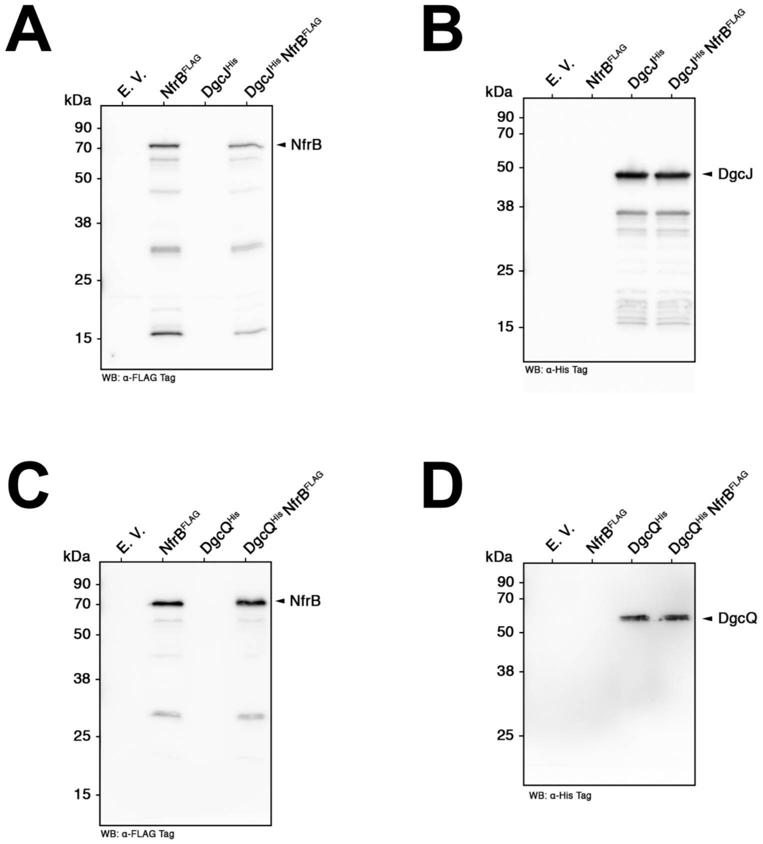


**Figure S3. Relevant protein levels in cellular samples used for the co-purification experiment shown in Fig. 5.** Protein levels were analyzed by immunoblot analysis for DgcJ$^{His}$ (A), NfrB$^{FLAG}$ (B, D) and DgcQ$^{His}$ (C) before the cell lysis procedure was applied.

### Supplementary Table

### Table S1

#### Oligonucleotide primers used in the present study[1].

#### I. Primers used for cloning *nfrB* into pGEX-6P-1 (Cytiva 58-9546-48):

| | |
|---|---|
| *NfrB*$^{451-745}$_BamHI_f | cat**ggatcc**atggttaaccgcatcgtgcag |
| *NfrB*$^{451-745}$_EcoRI_r | gta**gaattc**ttattctccttcattttcggactc |

#### II. Primers used for generating the *nfrB::lacZ* reporter gene fusion:

| | |
|---|---|
| *nfrBlacZ-1499_f* | catat**gaattc**gcatggatcatgatccactc |
| *nfrBlacZ+0_r* | gat**aagctt**ccatacaaaaaccttacattaacg |

#### III. Primers used for generating chromosomal knockout mutations by one-step inactivation (OSI):

| | |
|---|---|
| *nfrB*-H1P1 | gccaccctaaacataaccagcgttaatgtaaggttttgt**gtgtaggctggagctgcttc** |
| *nfrB*-H2P2 | aggttattctccttcattttcggactccagttgcgcaacc**attccggggatccgtcgacc** |
| *nfrA*-H1P1 | acaggttgcgcaactggagtccgaaaatgaaggagaataa**gtgtaggctggagctgcttc** |
| *nfrA*-H2P2 | gaaaatgaacttacgcatttaccagtgcactccaatggtg**attccggggatccgtcgacc** |
| *ybcH*-H1P1 | gcgcaacaacgcgtttctcaccattggagtgcactggtaa**gtgtaggctggagctgcttc** |
| *ybcH*-H2P2 | gaaagaggtaagccaggtcgtacccgacttacctggaggagat**attccggggatccgtcgacc** |
| *wecA-G*-H1P1 | ggtcttcgtggttatacttctgctaataattttctctgagagcatgcattgt**gtaggctggagctgcttc** |
| *wecA-G*-H2P2 | gcagacaggcgacggagtgaccactccgtcgctttacaaagagaggaaaa**attccggggatccgtcgacc** |
| *wecB*-H1P1 | gagcgcaaaggcgctcgccgcttattcgaagagaatcgat**gtggtgtaggctggagctgcttc** |
| *wecB*-H2P2 | gaaatggtcgcaaaactcatagtgatatccgattatttttaacgc**attccggggatccgtcgacc** |
| *wecC*-H1P1 | ctcgcattctggaagcgttaaaaaataatcggatatcactatga**gtgtaggctggagctgcttc** |
| *wecC*-H2P2 | gttatcagaatttttctcatcagcgccagactcctttggcatcgac**attccggggatccgtcgacc** |

#### IV. Primers used for generating the plasmid pAP58 by replacing the antibiotic resistance cassette of pCAB18[2] with the chloramphenicol resistance cassette of pACYC184[3]:

| | |
|---|---|
| pCAB18-BglII-rew | **ggcggc**agatct**gttgaatactcat**actcttcc |

---

[1] Relevant nucleotides (e.g. restriction sites, mutations introduced or pKD13-specific sequences) labeled in **bold**. All primer sequences are given from 5´- to 3´-end.

[2] Barembruch, C., and R. Hengge. 2007. Cellular levels and activity of the flagellar sigma factor FliA of *Escherichia coli* are controlled by FlgM-modulated proteolysis. Mol. Microbiol. 65:76-89.

[3] Chang, A. C. Y., and S. N. Cohen. 1978. Construction and characterization of amplifiable multicopy DNA cloning vehicles derived from the P15A cryptic miniplasmid. J. Bacteriol. 134:1141-1156.

| | |
|---|---|
| pCAB18-NdeI-for | ggcggc**catatg**ctgtcagaccaagtttactc |
| CAT-BglII-for | ggcggc**agatct**ggtgcttttgccgttacgcac |
| CAT-NdeI-rew | ggcggc**catatg**aataactgccttaaaaaaattac**gcc** |

#### V. Primers used for cloning *nfrBA-ybcH* and mutant alleles into pAP58:

| | |
|---|---|
| *nfrB*_XmaI_f | ct**cccggg**acataaccagcgttaatgtaag |
| *ybcH*_XbaI_r | ga**tctaga**ttaatactcgagaatgccgtg |
| *nfrB*$^{L490A}$_f | cgcccg**gcg**ggtcaaattc |
| *nfrB*$^{L490A}$_r | gaatttgacc**cgc**cgggcg |
| *nfrB*$^{L537A}$_f | gctggcacaggcg**gcg**gcagagcaaaac |
| *nfrB*$^{L537A}$_r | gttttgctctgc**cgc**cgcctgtgccagc |
| *nfrB*$^{G491L}$_f | gttgcgcccgtta**ctg**caaattctgctgg |
| *nfrB*$^{G491L}$_r | ccagcagaatttg**cag**taacgggcgcaac |
| *nfrB*$^{D169A}$_f | attctgcat**gcc**gccgaagatgtgatttc |
| *nfrB*$^{D169A}$_r | gaaatcacatcttcggc**ggc**atgcagaat |
| *nfrB*$^{D267A}$_f | gagtcttactgaa**gcg**tacgacattggcttc |
| *nfrB*$^{D267A}$_r | gaagccaatgtcgta**cgc**ttcagtaagactc |
| *nfrB*$^{W330A}$_f | atccc**gcg**cgatcatcggcattgttttc |
| *nfrB*$^{W330A}$_r | gaaaacaatgccgatgatcg**cgc**gggat |
| *nfrA*_BamHI_r | gta**ggatcc**gatcgtccagctg |
| *nfrB_ClaI_rev* | catt**atcgat**ggattcccacgcc |

#### VI. Primers used for cloning *dgcJ* and *dgcQ* into pRH800 (Lange and Hengge-Aronis, 1994) and generating mutant alleles:

| | |
|---|---|
| *dgcJ*_BamHI_f | cat**ggatcc**ctcgtttcactaaccgaagg |
| *dgcJ*_XbaI_r | gt**tctaga**tcatgaacggctgtttttgttc |
| *dgcJ*$^{6xHIS}$_XbaI_r | gt**tctaga**tcagtgatggtgatggtgatgtgaacggctgtttttgttc |
| *dgcJ*$^{GGAAF}$_f | ctcggtggc**gctgca**ttctgcatc |
| *dgcJ*$^{GGAAF}$_r | gatgcagaa**tgcagc**gccaccgag |
| *dgcQ*_BamHI_f | cata**ggatcc**gaatcataaaaaagcaggttggg |
| *dgcQ*$^{6xHIS}$_XbaI_r | at**tctaga**ttagtgatggtgatggtgatgagcgttatcgctcgcga |

#### VII. Primers used for cloning *nfrB*$^{3xFLAG}$ into pAP58:

| | |
|---|---|
| *NfrB*$^{3xFLAG}$*_f* | cacgacatcgactacaaggacgacgacgacaag**caactggagtccgaaaatg** |
| *NfrB*$^{3xFLAG}$*_r* | gtccttgtagtcaccgtcgtggtccttgtagtc**cgcaacctgttctgtgttta** |

#### VIII. Primers used for chromosomal C-terminal 3xFLAG-tagging of *nfrB* via two-step mutagenesis:

| | |
|---|---|
| *nfrB_ccdB_f* | tcgttcagcaattaacgtgttgttattgcgccatgaacgcagttctctgccgctcggcaa **cgcatcgtggccggatcttgc** |
| *nfrB_ccdB_r* | ggcgttctgccgcgcttaatgtgttctgctggcgtgcttcattgtatagcgtatctgcct **cggataacagaaaggccggg** |
| *nfrB*$^{3xFLAG}$*_f* | tgccgcatcagttcctgttc |
| *nfrB*$^{3xFLAG}$*_r* | cagtgccaggatccgatcgt |
